# FlashFold: a standalone command-line tool for accelerated protein structure and stoichiometry prediction

**DOI:** 10.1101/2025.01.23.634437

**Authors:** Chayan Kumar Saha, Mohammad Roghanian, Susanne Häussler, Lionel Guy

## Abstract

AlphaFold has revolutionized the decades-old issue of precisely predicting protein structures. However, its high accuracy relies on a computationally intensive step that involves searching vast databases for homologous sequences as the query protein of interest. Additionally, predicting the quaternary structure of protein complexes requires prior knowledge of subunit counts, a prerequisite rarely met. To address these limitations, we introduce FlashFold – a fast, user-friendly tool for protein structure prediction. It accelerates homology searches using a compact built-in database, enabling structure predictions up to 3-fold faster than AlphaFold3, with sacrificing little or no accuracy. Unlike others, FlashFold features adaptable built-in databases that allow users to easily incorporate their own privately sequenced data – an option that can positively influence the prediction accuracy. Moreover, it allows users to estimate stoichiometry of protein complexes directly from sequence information. To support high-throughput workflows and streamline downstream decision-making, it generates interactive and filterable summary reports, enabling users to efficiently visualize protein structures, and interpret large volumes of prediction results. FlashFold runs locally on Linux and Mac, eliminating reliance on third-party servers. FlashFold is available at https://github.com/chayan7/flashfold.

## Introduction

Proteins are responsible for all essential cellular functions and their tertiary structure is key to understanding their role. However, resolving protein structures using conventional experimental approaches like X-ray crystallography or Cryo-EM requires significant amount of time and effort, and thus researchers have long attempted to use *in silico* solutions instead. In 2020, DeepMind released AlphaFold2, an advanced machine-learning tool that predicts three-dimensional structures of single-chain proteins with unmatched accuracy, outperforming all competitors at 14th <u>C</u>ritical <u>A</u>ssessment of protein <u>S</u>tructure <u>P</u>rediction competition (CASP)^1^. AlphaFold-Multimer, a later extension of AlphaFold2, could accurately predict the structure of protein complexes in many cases, allowing researchers to explore protein-protein interactions (PPI) ^2^. To predict protein structures, AlphaFold relies on a novel end-to-end deep neural network that utilizes evolutionary information from multiple sequence alignments (MSA) of homologous sequences as its main input. Therefore, building reliable MSA is crucial, as the diversity of MSA influences the prediction accuracy ^1,3^.

For finding homologues as input, AlphaFold2 utilizes highly sensitive algorithms, HMMer ^4^ and HHblits ^5^, to search a comprehensive collection of protein sequences from publicly available resources and environmental databases. Due to the large data volume, 3 TB storage on a local computer is required, and MSA construction often takes hours, significantly impacting overall prediction time ^6^. Additionally, AlphaFold2 requires graphics processing units (GPUs) with a considerable quantity of GPU RAM (Random Access Memory) of 16 GB ^1,6^ to execute the deep neural network algorithm on query proteins of about 1000 residues.

To make protein structure prediction computationally more accessible and less expensive, ColabFold was released, achieving similar accuracy as AlphFold2 and AlphaFold-Multimer, but in a fraction of the time ^6,7^. Apart from speed, ColabFold also stands out for its lack of GPU limitations. However, the presence of a parallel computing platform-compatible GPU (e.g. CUDA 12.1 or later for ColabFold version 1.5.5) speeds up structure inference and energy minimisation. ColabFold reduces database searching time by using the more efficient, server-based MMseqs2 homology detection software to produce MSA ^6,8^. However, accessing the dedicated homology search server requires continuous internet connection. Tasks are also often standing in queue to start, due to the server offering only limited access for each user. To operate independently of the MMseqs2-based homology search server, users can download the smaller (1 TB) built-in ColabFold sequence database. However, the local ColabFold still requires high-performance computing, as the entire database must be loaded into RAM before homology detection begins, making the process time-consuming for single queries.

Recently, DeepMind released AlphaFold3 ^9^, the third major version of AlphaFold, which introduces the ability to predict complex structure of proteins, nucleic acids, small molecules, ions and modified residues. Initially AlphaFold3 was only accessible via a dedicated commercial server that limited users to a maximum of 20 job submissions per day ^9^, making it unsuitable for high throughput analyses and the pseudocode raised controversy ^10^. Eventually, the code was released, though not strictly in open-source fashion, since model weights – the core of the prediction software – are accessible only upon request. Nonetheless, users may now download and use the software for non-commercial purposes ^11^. It can only be installed on a Linux-based computer having at least 64 GB of RAM, an expensive NVIDIA GPU with Compute Capability 8.0, and about 1TB disk space (preferably SSD) for storing the sequence database. Moreover, the built-in databases offered by AlphaFold3, or its previous version AlphaFold2 or ColabFold cannot be modified, for example by adding newly sequenced or private data. Another challenge in existing methods is their reliance on prior stoichiometry information for protein complex structure prediction, which is often unknown. It raises the need for sequence-based stoichiometry prediction – a largely unexplored challenge with a few existing solutions ^12,13^.

Given the limitations of the currently available methods discussed above, our aim was to develop a tool that can be run locally, using a compact database, allowing users to extend it with additional sequence from public or private resources, which might enrich the MSA information content. MSA enrichment has been shown to be a successful strategy, significantly improving the prediction accuracy of ColabFold ^14^. Here, we present FlashFold, a python-based tool that; i) predicts the structure of protein monomers and multimers with an accuracy comparable to AlphaFold3, ii) predicts the subunit composition of a protein complex of unknown stoichiometry information, iii) provides a database at least an order of magnitude smaller than the ones offered by AlphaFold2, AlphaFold3, and ColabFold, and allows to build custom databases from public or private sequences, iv) can be installed on local computer or cluster running Linux or Mac operating system, and is independent of any third-party server, and v) reduces GPU limitations and ensures proper utilization of computer resources by keeping CPU and GPU based calculation separated.

## Results

### 1. Structure prediction

Protein structure prediction using FlashFold begins with a sensitive homology search to build a diverse MSA for three-dimensional structure prediction (Fig. 1). Upon completing structure prediction, FlashFold automatically generates a summary table containing key scoring metrics for the predicted models. Moreover, to estimate the quality of each interface on protein complexes, it calculates the pDockQ2 score ^15^.

**Figure 1.**
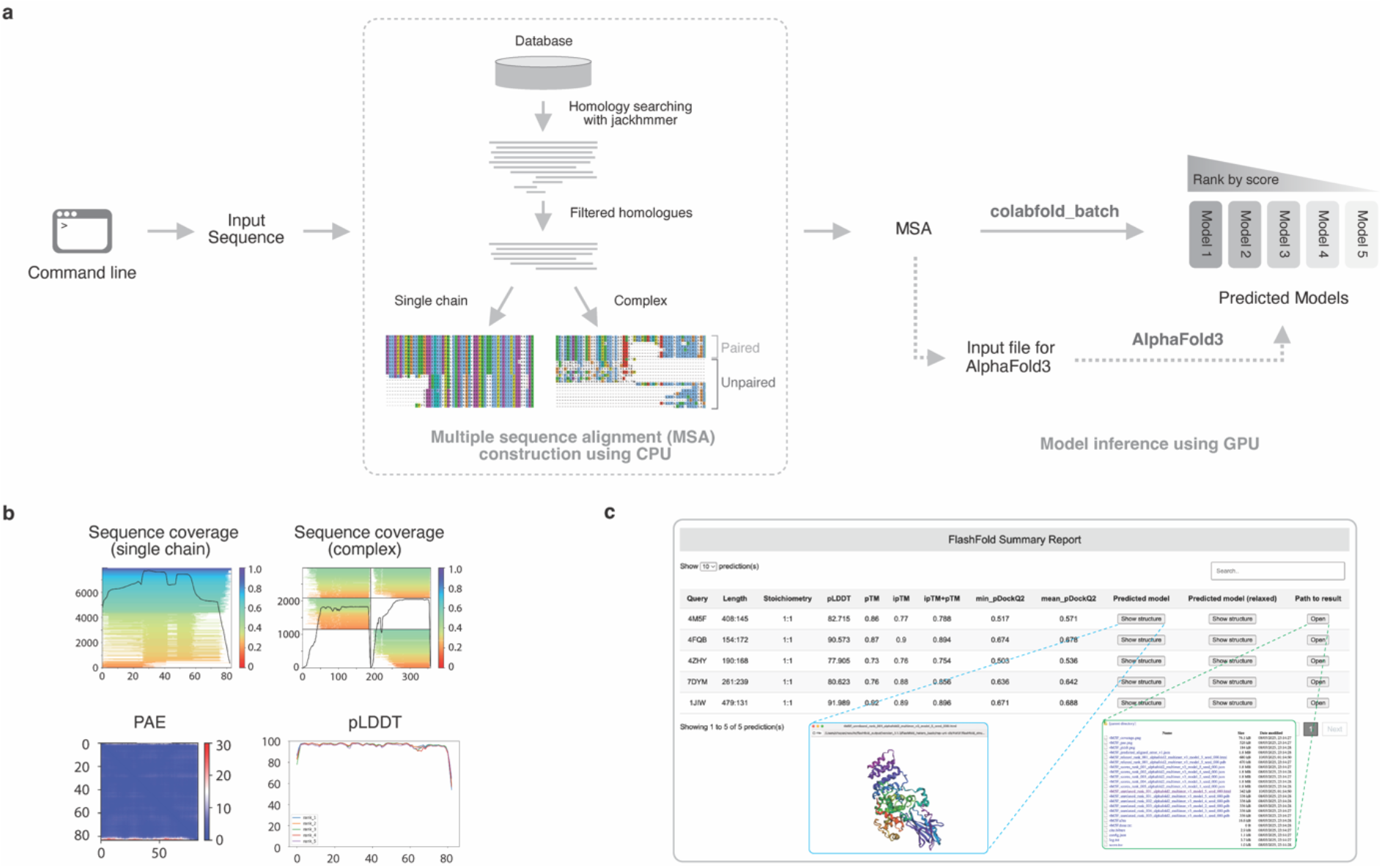
Overview of FlashFold. **(a)** FlashFold uses FASTA sequences as input to search for related sequences against a very compact built-in sequence database using Jackhmmer ^4^. Homologous sequences are filtered and processed as an MSA file in A3M format, and then sent to colabfold_batch for structure prediction. For single chain prediction, only non-redundant homologous sequences are chosen. FlashFold can optionally employ a strict filtering step to create a compact MSA by clustering the detected homologues using CD-hit tool and selecting representative sequences from the clusters. For complex structure prediction, the final MSA is the combination of paired and unpaired MSA. Jackhmmer hits with the lowest e-values from the same species are paired and only non-redundant paired sequences are selected. Unpaired MSA is produced using the same strategy as for single chain predictions, with gaps in place of missing homologues. FlashFold optionally offers the user to compute structure inference using AlphaFold3. Predicted models are ranked by decreasing quality score. (**b**) FlashFold output includes figures depicting MSA depth and coverage, and AlphaFold2 confidence metrics such as PAE and pLDDT, to evaluate prediction quality. (**c**) FlashFold also compiles a table of model quality scores and interface attributes at the end of the prediction phase, formatted as a responsive HTML summary report.

Three compact sequence databases are included with FlashFold to facilitate quicker homology detection (Fig. 2a, see Method 2). To compare the structure prediction performance with AlphaFold2 and AlphaFold3, we implemented FlashFold to predict structure with a custom dataset of 15 monomeric and 10 multimeric proteins from *P. aeruginosa* (Supplement Table 1, Methods 7.1). The predictions could be then compared to the structures already experimentally determined and published in Protein Data Bank (PDB) ^16^. On average, FlashFold generated MSA considerably faster than AlphaFold (Fig. 2b), accelerating overall predictions by up to threefold and sixfold compared to AlphaFold3 and AlphaFold2, respectively (Fig. 2c), while maintaining reliable accuracy (Fig. 2d). Using a smaller database, such as *rep-bac-db*, significantly enhances MSA generation speed, making it six times faster than AlphaFold3 and fourteen times faster than AlphaFold2. To compare prediction accuracy, we calculated the TM-score using TM-align ^17^, which compares protein structure in a sequence independent manner. FlashFold achieved a mean TM-score of 0.90, compared to AlphaFold2 and AlphaFold3’s 0.91, indicating a “very high” similarity between the predicted and experimentally resolved structures (see predicted hetero-complexes in Fig. 3a, superimposed with the native structures using PyMol v2.5.4).

**Table 1.** Twenty-five randomly selected *Pseudomonas aeruginosa* proteins and complexes used for benchmarking. The first 15 single chain proteins were considered for run time benchmarking but all of them were included for the prediction accuracy benchmark.

| Serial | PDB ID | UniProt Accession | Gene Name | Oligomeric state |
| --- | --- | --- | --- | --- |
| 1 | 1azu | P00282 | *azu* | Monomer |
| 2 | 1ddk | Q79MP6 | *blaIMP-1* | Monomer |
| 3 | 1dvv | P00099 | *nirM* | Monomer |
| 4 | 1ex9 | P26876 | *lip* | Monomer |
| 5 | 1g68 | P16897 | *pse4* | Monomer |
| 6 | 2fgo | Q9I6D2 | *fdx1* | Monomer |
| 7 | 2kep | Q00514 | *xcpT* | Monomer |
| 8 | 2lc4 | Q51354 | *pilP* | Monomer |
| 9 | 2lhf | Q51486 | *oprH* | Monomer |
| 10 | 2x27 | Q9HWW1 | *oprG* | Monomer |
| 11 | 3jyz | Q8KQ36 | *pilA* (Pilin) | Monomer |
| 12 | 4nx9 | O86266 | *fliC* | Monomer |
| 13 | 4rlc | P13794 | *oprF* | Monomer |
| 14 | 6bbk | P18774 | *pilA* (Fimbrial protein) | Monomer |
| 15 | 6fip | Q51368 | *tonB* | Monomer |
| 16 | 1bex | P00282 | *azu* | Homodimer |
| 17 | 1fof | P14489 | *bla* | Homodimer |
| 18 | 1ix1 | Q9I7A8 | *def* | Homodimer |
| 19 | 6p90 | Q9I4W3 | *dapA* | Homodimer |
| 20 | 7jrk | O68562 | *bamE* | Homodimer |
| 21 | 1jiw | Q03023 - Q03026 | *aprA - inh* | Heterodimer |
| 22 | 4fqb | Q9I2Q - Q9I2Q0 | *tse1 - tsi1* | Heterodimer |
| 23 | 4m5f | Q9HYC5 - Q9HYC4 | *tse3 - tsi3* | Heterodimer |
| 24 | 4zhy | Q9I4L4 - Q9I4L6 | *yfiR - yfiB* | Heterodimer |
| 25 | 7dym | Q9HXA6 - Q9HXA5 | *PA3907 - PA3908* | Heterodimer |

**Figure 2.**
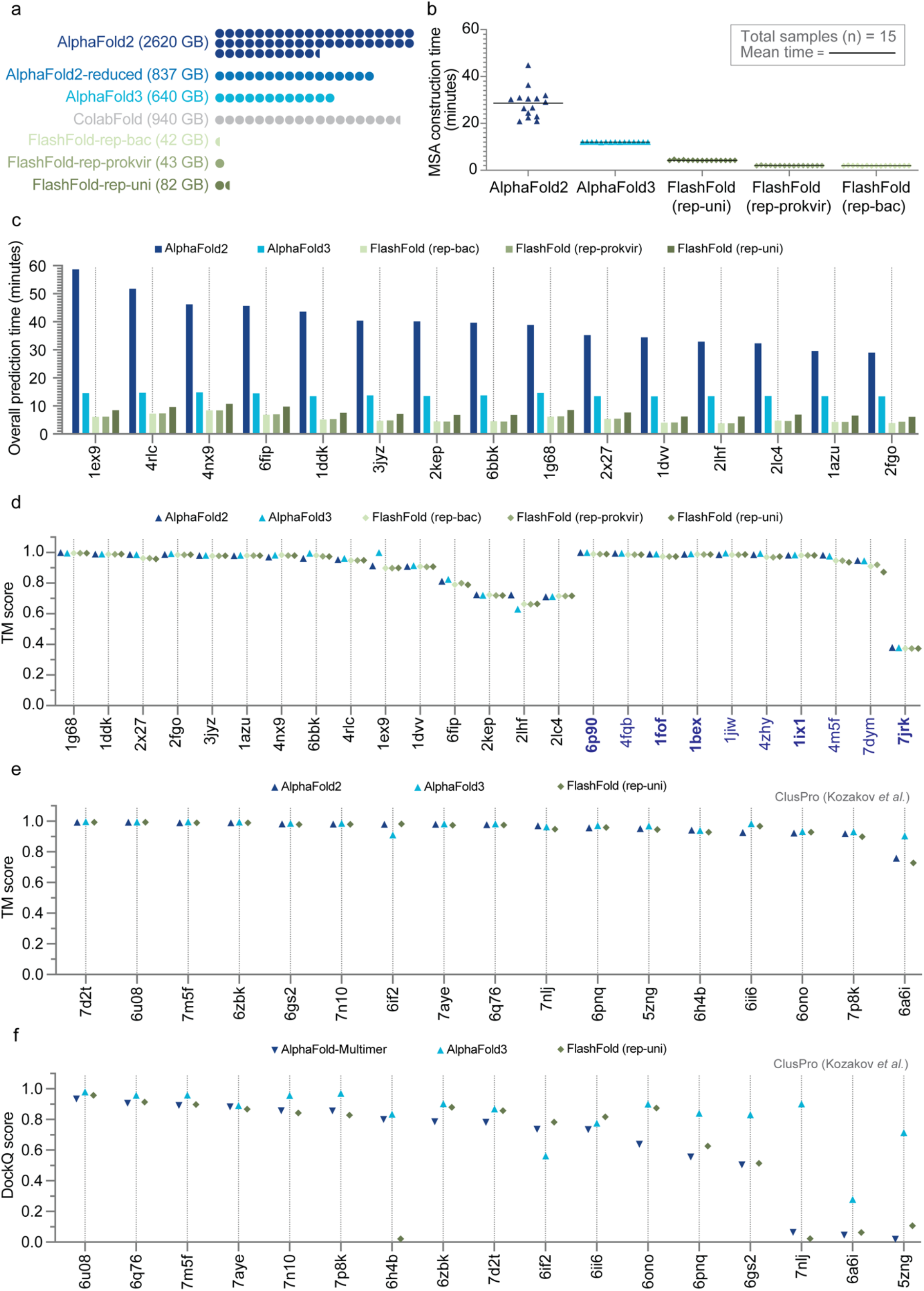
Performance of FlashFold. (**a**) FlashFold offers three built-in databases which are extremely compact compared to the databases offered by AlphaFold2, AlphaFold3 and ColabFold. The largest of these three FlashFold databases, *rep_uni_db*, is generally applicable, while *rep_prokvir_db* is designed for prokaryotes and viruses, and *rep_bac_db* for bacteria only. (**b**) A comparison of MSA construction times for AlphaFold2, Alphafold3 and FlashFold. (**c**) A comparison of monomer structures prediction times for AlphaFold2, Alphafold3 and FlashFold. (**d**) Comparison of AlphaFold2, AlphaFold3 and FlashFold prediction accuracy for monomers and dimer complexes (coloured in blue and homodimers in bold). (**e**) Evaluating the accuracy of multimeric structure prediction of AlphaFold2, AlphaFold3 and FlashFold using ClusPro dataset. (**f**) Comparison of AlphaFold-Multimer, AlphaFold3 and FlashFold multimeric prediction modes on the ClusPro dataset. A reliability metric for protein-protein docking models known as DockQ was calculated and compared.

**Figure 3.**
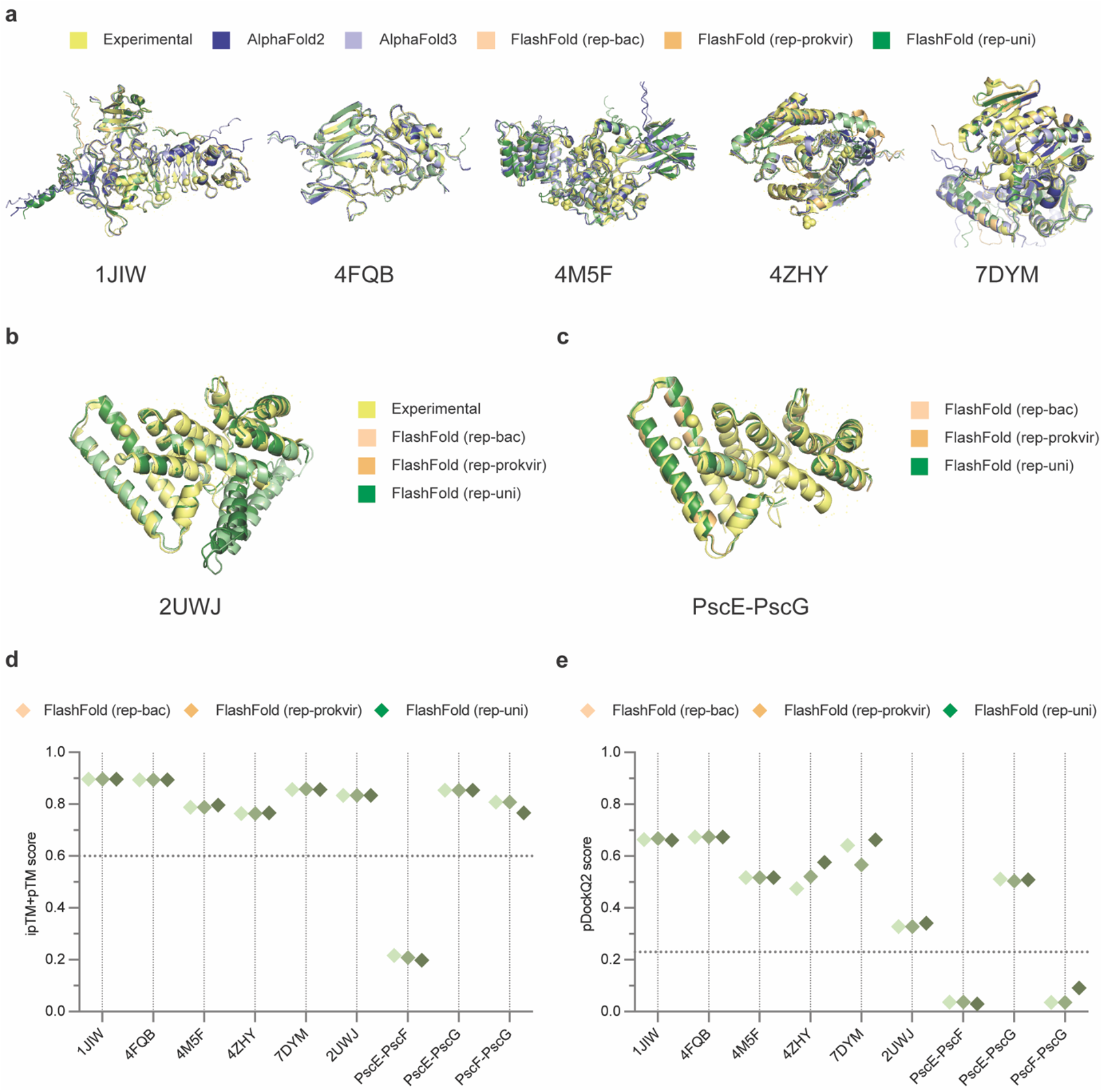
FlashFold can be used to differentiate between interacting protein partners from the noninteracting ones. 1JIW, 4FQB, 4M5F, 7DYM and 4ZHY are known heterodimer complex PDB IDs from *Pseudomonas aeruginosa* (Table 1). 2UWJ is a PDB ID of a ternary (1:1:1) complex that includes PscE, PscF and PscG proteins. PscF is a primary part of the *P. aeruginosa* type three secretion needle, and researchers found that using gel-filtration techniques, a complex of its other two cytoplasmic partners, PscE and PscG, could be separated, suggesting a direct interaction between these two macromolecules ^26^. Based on FlashFold prediction across its three built-in databases, (**a**) predicted models of 1JIW, 4FQB, 4M5F, 7DYM, 4ZHY, (**b**) PscE-PscF-PscG complex (2UWJ), and surprisingly (**c**) PscE-PscG complex (which lacks an experimentally resolved structure) all cleared the FlashFold PPI threshold (Methods 6) with the required predicted pTM score (**d**) and pDockQ2 score (**e**). It suggests that the FlashFold PPI threshold could identify protein pairs that do not directly interact such as PscE-PscF and PscF-PscG.

Additionally, we used a dataset from ClusPro ^18^ – a web server dedicated to predicting models of the complex using the structures of interacting proteins – to benchmark the multimeric prediction accuracy of FlashFold (Methods 7.2). FlashFold nearly always achieved TM-scores that were equivalent to those of AlphaFold2 and AlphaFold3 (Fig. 2e). Moreover, the predicted DockQ scores, a reliability metric for protein-protein docking models ^19^, were nearly always better with FlashFold when compared to AlphaFold-Multimer (Fig. 2f) ^2,18^. For analysing the ClusPro dataset, we only used the FlashFold *rep_uni_db*, because the dataset contains query protein sequence from eukaryotes.

Another unique feature of the FlashFold built-in databases is that they can be complemented by sequences from both public and private resources, e.g. from a library of genomes from a species of interest. The inclusion of additional sequence data has the potential to enrich the MSA with diverse evolutionary signals, which was previously reported to contribute to improved prediction accuracy ^14^. We expanded the FlashFold built-in database using sequence data from *P. aeruginosa* that is publicly available in the NCBI RefSeq, as well as some from private resources (Methods 7.3). We found this additional evolutionary information led to an increased TM-score for most test cases (Table 2).

**Table 2.**
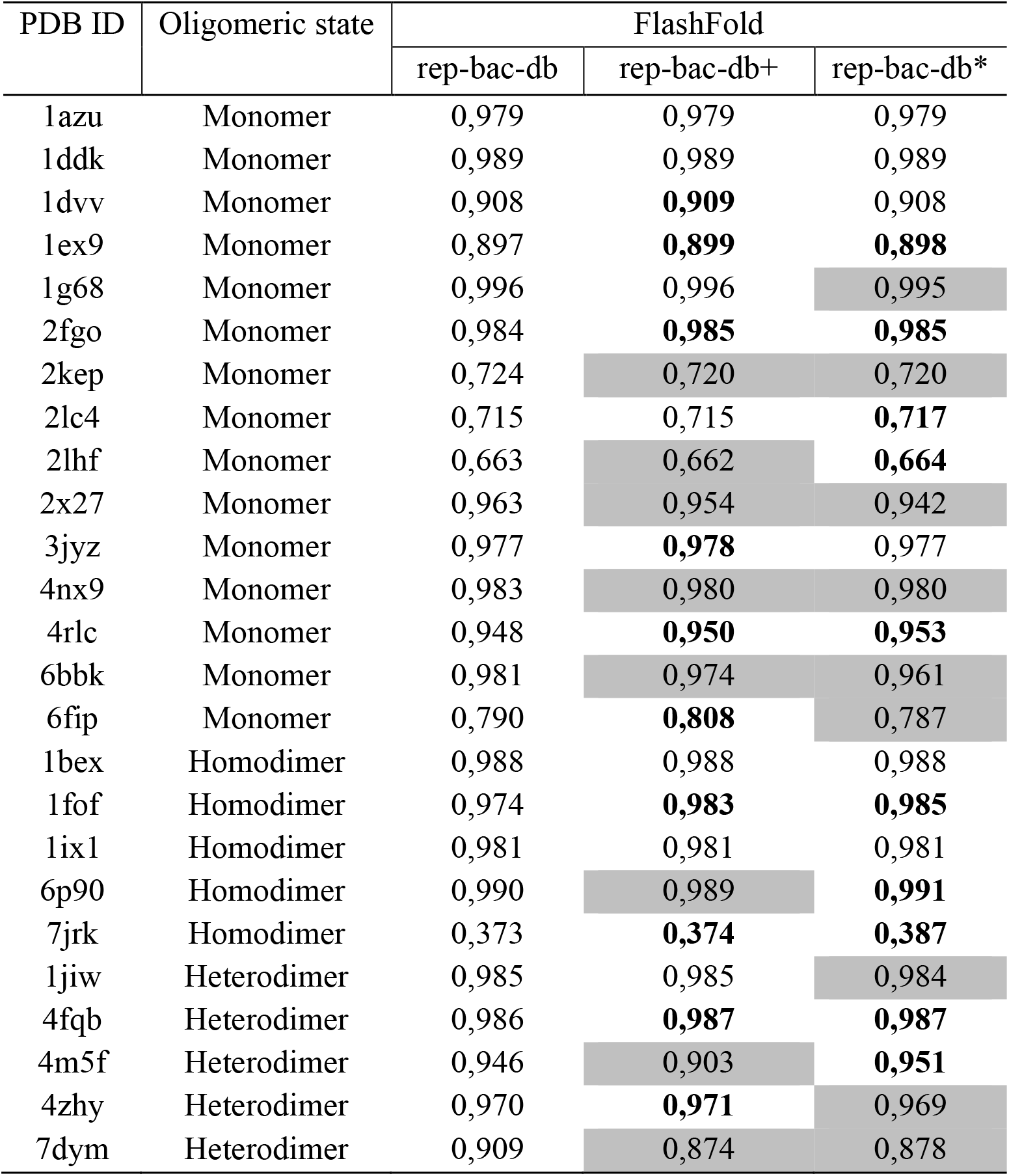
FlashFold database extension with sequence from public and private resources impacts prediction accuracy. We used twenty-five *P. aeruginosa* proteins and complexes (Table 1) for this analysis. The TM scores were influenced in many cases (increasing scores are shown as bold and decreasing scores are shaded grey) when we used FlashFold with *rep-bac-db+* or *rep-bac-db\** instead of *rep-bac-db*.

### 2. Stoichiometry prediction

We also assessed the applicability of FlashFold in predicting the stoichiometry of protein complexes. For this analysis, we selected nine queries – comprising three monomers, three homodimers, and three heterodimers – from the customized benchmarking dataset (Table 1). In eight of the nine tested cases, using only sequence information of the query input, FlashFold accurately predicted the stoichiometry as that was indicated in the publicly available experimentally determined structure (Table 3). It failed to detect correct oligomeric state for UniProt accession O68562 (shaded grey in the Table 3). According to PDB database, it is a homodimer (A2), but FlashFold predicted it as a monomer (A1). We did not employ AlphaFold3 to benchmark stoichiometry prediction; nevertheless, FlashFold optionally allows users to integrate AlphaFold3.

**Table 3.** Predicting stoichiometry using FlashFold. We examined nine *P. aeruginosa* proteins and complexes for this analysis comprising three monomers, three homodimers, and three heterodimers. In almost every case except one (UniProt accession O68562, shaded in grey), FlashFold correctly predicted the stoichiometry.

| Serial | UniProt accession | PDB ID | PDB annotated stoichiometry | Tested stoichiometry combinations | Total tested stoichiometry combinations | FlashFold predicted stoichiometry |
| --- | --- | --- | --- | --- | --- | --- |
| 1 | P00282 | 1azu | A1 | A1, A2, A3 | 3 | A1 |
| 2 | Q79MP6 | 1ddk | A1 | A1, A2, A3 | 3 | A1 |
| 3 | P00099 | 1dvv | A1 | A1, A2, A3 | 3 | A1 |
| 4 | P14489 | 1fof | A2 | A1, A2, A3 | 3 | A2 |
| 5 | Q9I4W3 | 6p90 | A2 | A1, A2, A3 | 3 | A2 |
| 6 | O68562 | 7jrk | A2 | A1, A2, A3 | 3 | A1 |
| 7 | Q03023-Q03026 | 1jiw | A1B1 | A1B1, A1B2,<br>A1B3, A2B1,<br>A2B2, A2B3,<br>A3B1, A3B2,<br>A3B3 | 9 | A1B1 |
| 8 | Q9HYC5-Q9HYC4 | 4m5f | A1B1 | A1B1, A1B2,<br>A1B3, A2B1,<br>A2B2, A2B3,<br>A3B1, A3B2,<br>A3B3 | 9 | A1B1 |
| 9 | Q9I4L4-Q9I4L6 | 4zhy | A1B1 | A1B1, A1B2,<br>A1B3, A2B1,<br>A2B2, A2B3,<br>A3B1, A3B2,<br>A3B3 | 9 | A1B1 |

### 3. Predict protein-protein interaction

Lastly, we evaluated the potential of FlashFold to predict novel PPIs. By default, as a part of structure prediction of protein complexes, FlashFold generates a confidence score and calculates pDockQ2 score for each predicted model. We chose a model confidence cut-off score of 0.6 and a pDockQ2 cut-off score of 0.23 for PPI prediction (see details in Methods 6). Using the threshold, we revalidated seven previously identified interactions, six of which have resolved structures (Fig. 3a-c). Furthermore, FlashFold could effectively distinguish between interacting and non-interacting protein partners (Fig. 3d and 3e), highlighting its potential for large-scale identification of novel PPIs.

## Discussion

In this contribution, we present FlashFold, a standalone tool for rapid and accurate structure prediction. Our benchmarking results demonstrate that FlashFold significantly surpasses state-of-the-art methods in speed while maintaining reliable prediction accuracy. FlashFold optimizes computational resource utilization and ensures efficient performance by decoupling the CPU-intensive MSA generation from the GPU-dependent structure inference. Moreover, users can easily elect to use AlphaFold3 for structure inference – a valuable option given that AlphaFold3 outperforms AlphaFold2 on easier targets, though not consistently on more challenging ones, according to CASP rankings ^20^. FlashFold allows users to strategically select the most appropriate inference model per case, making it more adaptable. We believe that FlashFold stands out from other tools with its compact, built-in sequence database, offering a unique blend of speed, efficiency, and accuracy.

Unlike other tools, FlashFold empowers users with a dedicated script to build as well as modify sequence database using both public and private sequence data. Its structured database ensures full traceability of sequence co-evolutionary information, a critical factor for generating paired MSAs essential for accurate complex structure prediction. Furthermore, the non-redundant and compact design of the database enhances its efficiency, enabling faster MSA construction (Fig. 2b and Fig. 2c) – a key bottleneck in AlphaFold2 and AlphaFold3. Most importantly, structure prediction using FlashFold is completely template independent, solely relies on the efficiency of sequence database in building a diverse MSA.

Our benchmark analysis, utilizing both the customized and ClusPro datasets, demonstrated its efficiency, as reflected in the prediction accuracy (Fig. 2d-2e). We also observed that addition of private sequence data to the in-built sequence database can increase the prediction accuracy, highlighting the significance of the modifiable feature of FlashFold database (Table 2).

FlashFold has a great potential for uncovering novel PPIs on a high-throughput scale. For complex models, it generates a score-based table as the final step of prediction, providing a quantitative reference for PPI assessment. Our analyses indicate that these scores effectively distinguish interacting protein partners from non-interacting ones (Fig. 3).

Another potential application for FlashFold is predicting the stoichiometry of protein complexes, using only sequence information. In our analyses, it could predict stoichiometry of eight out of nine complexes accurately (Table 3). In addition to these, FlashFold offers user to modify the MSA and add ligand before structure prediction. To utilize this feature, user needs to have AlphaFold3 installed in their local machine.

In summary, FlashFold offers a comprehensive solution for protein structure prediction on a high-throughput scale. Furthermore, it generates interactive and filterable summary reports to visualize results in a structured manner. For researchers with access to private sequencing data, it serves as a valuable resource for exploring how additional sequence information may influence the structure of their protein of interest or its possible interactions.

## Methods

### 1. Implementing FlashFold

FlashFold is a standalone user-friendly program, with an emphasis on being computationally less demanding so that researchers can predict structures of proteins of interest faster on a high-throughput manner using their local machine. It can be installed on Linux or Mac operating system from its source at https://github.com/chayan7/flashfold. It takes protein sequences in FASTA files as the primary input.

Structure prediction using FlashFold consists of three main steps (Fig. 1). First, a sensitive homology detection is performed using the Hidden Markov Model-based method Jackhmmer, part of the HMMer distribution ^4^, to construct a MSA (Methods 3). Second, the MSA is used as an input for the structure prediction using the colabfold_batch script, part of the ColabFold package ^6^. Third, a summary table is generated containing the scoring matrices for the predicted structures, including pDockQ2 score for protein complexes which estimates the quality of each interface in a multimer model ^15^.

Moreover, FlashFold allows user to predict structures using the FlashFold-generated MSA or user-specified custom MSA file in A3M format. In the latter case, the CPU-dependent homology detection step will be skipped. FlashFold also offers a batch prediction mode (“--batch” option), with which the user can input a directory that contains files formatted as FASTA or A3M.

FlashFold provides flexibility in the output generated, e.g. to only generate MSA (using the “--only_msa” option), formatting it to a JSON file to be used as input for AlphaFold3 (using the “--only_json” option). Users may also then integrate ligand information in the JSON file (using the “ligand” subcommand). Finally, if AlphaFold3 is installed locally alongside FlashFold, users can also perform structure prediction with it, using the “run_af3” subcommand.

### 2. FlashFold compact databases

To execute AlphaFold2, AlphaFold3 or ColabFold in a local machine, the user needs to download the databases which require a very large storage space (Fig 2a). Unlike these, FlashFold built-in databases are extremely compact. Using publicly available sequence information obtained from NCBI RefSeq ^21^, we created three unique databases for different application.

First, we built a database specific for bacterial proteins. The database is called *rep_bac_db*, (“<u>rep</u>resentative <u>ba</u>cterial <u>d</u>ata<u>b</u>ase”). To build it, we downloaded GenBank files of all the representative bacterial genomes using FlashFold subcommand *ncbi_data*. Then, to create a database from the downloaded GenBank files, we used the FlashFold subcommand *create_db*. It generates a FASTA file named “sequence_db.fasta” containing unique protein sequences extracted from all GenBank files, which is then used as a database for homology detection. Each protein sequence is assigned to a unique hash value, which is used as a key for accession sequence related information. To determine which taxa the protein belongs to, a file called “protein_to_gbks.json” is being generated, which contains a list of taxa (as hash value) for each sequence hash key. It also generates another file called “prot_hash_to_accession.json” to store a list of protein accessions against each protein hash key. The *rep_bac_db* database is 42 GB in size and contains 76,005,099 protein sequences from 19,695 bacterial species.

Second, we created a *rep_prokvir_db*, short for “<u>rep</u>resentative <u>prok</u>aryotic and <u>vir</u>al <u>d</u>ata<u>b</u>ase” which is an extended version of *rep_bac_db* with additional sequence information retrieved from archaea and viruses. We downloaded GenBank files of all the reference archaeal genomes. For viruses, we downloaded GenBank files from all the RefSeq annotated reference genomes. Then we extended the *rep_bac_db* using FlashFold subcommand *extend_db*. The rep_prok_db is 43 GB, contains 78,630,657 protein sequences from 35,220 taxa.

Third, we extended *rep_prokvir_db* with the sequence information from eukaryotes and named it as *rep_uni_db*, short for “<u>rep</u>resentative <u>uni</u>versal <u>d</u>ata<u>b</u>ase”. As previously, we downloaded GenBank files of all the RefSeq annotated reference eukaryotic genomes and later used FlashFold subcommand *extend_db* for extension. This is the largest database among all three (82 GB; Fig. 2a), contains 124,790,329 protein sequences from 37,244 taxa.

The user can download these abovementioned built-in databases using FlashFold subcommand *download_db* and apply any of these based on their research interest. The initial databases correspond to NCBI RefSeq database release 226 (September 13, 2024). They will be regularly updated, following the bimonthly updating schedule of RefSeq; however, user can also independently create and maintain these databases using FlashFold database-related subcommands.

The ability for the user to update the database as needed is a unique characteristic of FlashFold databases. Adding more sequences can influence the homology detection and the processing time of the MSA (Fig. 2b), but it can be useful to discover novel structural folds or PPI. FlashFold built-in databases are designed with an aim to make faster detection of distant homologues of the raw input sequence. It is also possible to enrich the MSA with closely related homologues by adding more information to the database. Below (Methods 7.3), we explain how we extended the *rep_bac_db* with additional sequence information from both public and private resources and checked its influence on prediction accuracy (Table 2).

### 3. Faster MSA construction with Jackhmmer

The compact nature of FlashFold databases enables a much faster, yet sensitive homology search. For this, FlashFold relies on Jackhmmer with the following parameters: “--F1 0.0005 --F2 0.00005 --F3 0.0000005 --incE 0.0001 -E 0.0001 -N 1”. For each input sequence, it performs a single iteration of alignment against the database (sequence_db.fasta file) by applying three filtering stages (--F1, --F2, and --F3) to progressively refine the search and finally accepts hits with an E-value ≤ 0.0001.

For single chain structure prediction, only non-redundant homologous sequences are selected to be included in the final MSA file. In addition, FlashFold offers an optional second filtering step (option “--compact_msa”) to create a more compact and diverse MSA. This step involves clustering the Jackhmmer detected homologues using CD-hit ^22^, with three cut-off values for sequence identity (70%, 80%, and 90%; with “-c” option). FlashFold selects the cut-off value that yields a number of clusters closest to 1000 and, a representative sequence from each cluster is then considered for MSA. Compact MSA is useful to speed up structure inference step, however it might result in reduced prediction accuracy. The MSA for homo-oligomeric complexes is constructed in the same way as the single chain, but the MSA is duplicated repeatedly for each component in the complex. For hetero-oligomeric complex structure prediction, FlashFold executes parallel homology searches for each component, using the same parameters mentioned before. It then creates one paired alignment and one unpaired alignment. To create paired MSA, firstly, for each species, it collects the best hits from each component and pairs them if the paired alignment covers at least 50% of the concatenated query sequence. Finally, from all paired alignment, it selects only the non-redundant ones. Unpaired MSA is created in the same way as described for monomer, with gaps in place of missing homologues for component.

### 4. Structure prediction

For structure prediction, FlashFold relies on the colabfold_batch script that comes along with ColabFold python package ^6^. The custom alignment (A3M format) produced by FlashFold is directly used as an input for colabfold_batch script, which mainly relies on AlphaFold2 models for structure inference. Moreover, FlashFold optionally allows the integration of AlphaFold3 for structure prediction. For single chain structure prediction, the pLDDT score is used to rank the models. It is a per residue measure of local confidence that spans from 1 to 100, with a higher score indicating a more accurate prediction ^1^. For complexes, it uses the predicted TM-score which is an integrated assessment of the overall predicted structure of the complex ^1^. To further evaluate the prediction quality, the output also includes graphics – generated by colabfold_batch – that show the sequence coverage in the MSA and AlphaFold2 confidence measures as in PAE that is also useful to rank and judge the quality of the predicted structures^1,6^.

### 5. Stoichiometry analysis

FlashFold comes with a dedicated subcommand, “stoi”, to predict the stoichiometry of a protein complex using only sequence information. Users can input a FASTA file with the sequences of the subunits of the query protein complex. For instance, to predict the stoichiometry of a complex with two subunits, A and B, the FASTA file should include their respective amino acid sequences.

Users can specify copy numbers for each subunit using the “--specific_stoichiometry” or “--global_stoichiometry” arguments. For example, if the copy number for both subunits A and B is set to 2 (using “--global_stoichiometry”), FlashFold generates four possible stoichiometric combinations (A1B1, A1B2, A2B1, A2B2) and predicts structures for each. It is also possible to assign different copy number for different subunits using “--specific_stoichiometry” option. Moreover, user can optionally integrate AlphaFold3 for the structure inference step during stoichiometry prediction.

The predicted models are ranked based on their prediction quality scores – interface predicted template modelling (ipTM) score for heterocomplexes and predicted template modelling (pTM) score for homomers. The stoichiometry combination with the highest score is considered the most likely oligomeric state. Additionally, FlashFold generates an interactive HTML summary report, allowing users to visualize all evaluated stoichiometries along with their corresponding quality scores. Using this approach, we predicted the stoichiometry of 9 different queries of which 3 are monomeric, and 6 were multimers (see Table 3).

### 6. Predict PPI using the FlashFold scoring matrices

FlashFold generates a scoring table for all the predicted models, ranked by prediction accuracy. In addition, for complexes, it calculates pDockQ2 score (script available from https://gitlab.com/ElofssonLab/huintaf2/-/blob/main/bin/pDockQ2.py) that estimates the quality of each interface in a multimer model ^15^. Protein complex models with a predicted TM-score > 0.6 are generally considered of high quality ^23^. Researchers have used predicted TM-score alone, or along with pDockQ2 score to predict interaction between subunits within a predicted multimer model ^15,23,24^. Protein complexes are deemed reliable when the pDockQ2 score exceeds 0.23 ^15^.

To guide PPI prediction, FlashFold calculates a model confidence score as shown below using ipTM and pTM score:

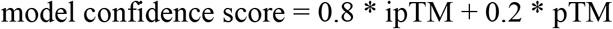

Moreover, FlashFold calculates pDockQ2 score and summarizes all the scores in a single table, offering the opportunity to confirm known, and discover novel, protein-protein interaction. We used the following criteria to identify seven previously known interactions (Fig. 3), using:

1. For heterodimers, the predicted model should have a model confidence score > 0.6 and the minimum of the pDockQ2 scores calculated for two chains should be > 0.23
2. For complexes that are not heterodimer, the predicted model should have a model confidence score > 0.6 and the mean of the pDockQ2 scores calculated for each chain should be > 0.23.

### 7. Benchmark

We utilised multiple datasets to benchmark FlashFold with AlphaFold2 and AlphaFold3, measuring both computing resources and levels of prediction accuracy. We used a Linux computer with Intel® Xeon® W-2295 18-core CPU, 64GB RAM and a NVIDIA GeForce RTX 3060 GPU with 12GB RAM. No template was used for FlashFold-based structure prediction. We have not considered benchmarking with ColabFold. In the case of ColabFold, the sequence search is server-based and should be run in a local machine with at least 768GB to 1TB RAM ^6,9^.

#### 7.1 Benchmark with a customised dataset of twenty-five *Pseudomonas aeruginosa* proteins and complexes

We randomly selected 25 proteins from *Pseudomonas aeruginosa* (Table 1) whose structures were already experimentally determined and published in Protein Data Bank (PDB) ^16^. We selected five homodimers, five heterodimers and 15 monomers. We extracted the sequences from UniProtKB Database and used them as input for structure prediction ^25^. For a fair comparison of MSA construction and overall run time, only monomer structure prediction data were included (Fig 2b and 2c), and both FlashFold and AlphaFold used the same number of CPUs. We calculated TM-score, a sequence independent measure of structural similarity, for each predicted models against the experimental structures using TM-align ^17^ and used it for comparing the prediction accuracy (Fig 2d).

#### 7.2 Benchmark with ClusPro dataset

We used both TM-align ^17^ and DockQ ^19^ to compare 17 ClusPro target predictions to their native structures ^2,18^. The predicted models from FlashFold were ranked by predicted interface TM-score (default) as returned by AlphaFold-Multimer. TM-scores were calculated for each highest ranked predicted model as mentioned in Methods 7.1 and was utilized to compare the prediction accuracy between AlphaFold and FlashFold (Fig 2e). The reference DockQ scores for the AlphaFold-Multimer were gathered from the Mirdita, et al. ^6^ (Fig 2f) and for rest were calculated with the DockQ tool using the native and highest ranked predicted models as input.

#### 7.3 Benchmark with extended FlashFold database

For this, we used the 25 previously selected proteins (Table 1) from *P. aeruginosa* for structure prediction. We extended the *rep-bac-db* with publicly and privately available *P. aeruginosa* specific sequence information to investigate whether the addition of new pathogen specific data influences the prediction accuracy. Consequently, we started by adding all the *P. aeruginosa* GenBank files that are publicly accessible in the NCBI RefSeq database (a total of 100,53 as of October 21, 2024) to the *rep-bac-db* and renamed the database to *rep-bac-db+* which is 46 GB in size and contains 1,822,441 additional unique protein sequences. Furthermore, we extended *rep-bac-db+* to *rep-bac-db\** by adding privately accessible GenBank files from a huge collection of clinical *P. aeruginosa* isolates (a total of 2,500 as of June 12, 2024, from Copenhagen Hospital). The *rep-bac-db\** has 215,438 additional unique sequences than that of *rep-bac-db+*, and a size of 47 GB. We later used these extended databases to predict structure. For each predicted model, we calculated the TM-score as mentioned in Methods 7.1 and compared the scores with that of *rep-bac-db* (Table 2).

## Data availability

The three in-built databases of FlashFold, namely *rep-bac-db, rep_prokvir_db* and *rep_uni_db* are readily available for download from: https://doi.org/10.5281/zenodo.14717354, https://doi.org/10.5281/zenodo.14720530, and https://doi.org/10.5281/zenodo.14720238 respectively. All the structures predicted during benchmark are available at: https://doi.org/10.5281/zenodo.15345646. The data used for generating the plots are available at https://doi.org/10.5281/zenodo.15345525.

## Code availability

FlashFold code is freely available under the MIT license at https://github.com/chayan7/flashfold.

The authors have no conflicting interests.

## Author contributions

All authors conceptualised and coordinated the study. C.K.S. developed the software and drafted the manuscript with contributions from all authors.

## Ethics declarations

